# Structure-specific regulation of nutrient absorption, metabolism and transfer in arbuscular mycorrhizal fungi

**DOI:** 10.1101/491811

**Authors:** Hiromu Kameoka, Taro Maeda, Nao Okuma, Masayoshi Kawaguchi

**Affiliations:** Division of Symbiotic Systems, National Institute for Basic Biology, 38 Nishigonaka, Myodaiji, Okazaki, Aichi 444-8585, Japan.; Graduate School of Life and Environmental Sciences, Osaka Prefecture University, 1-1 Gakuen-cho, Naka-ku, Sakai, Osaka 599-8531, Japan; The Graduate University for Advanced Studies, 38 Nishigonaka, Myodaiji, Okazaki, Aichi 444-8585, Japan.

**Author notes:** **Corresponding authors:** Hiromu Kameoka; Email address, Masayoshi Kawaguchi.

**Keywords:** Arbuscular mycorrhizal fungi, metabolism, nutrient absorption, nutrient transfer, transcriptome

## Abstract

Arbuscular mycorrhizal fungi (AMF) establish symbiotic relationships with most land plants, mainly for the purpose of nutrient exchange. Many studies have revealed the regulation of absorption, metabolism, and transfer of nutrients in AMF and the genes involved in these processes. However, the spatial regulation of the genes among the structures comprising each developmental stage are not well understood. Here, we demonstrate the structure-specific transcriptome of the model AMF species, *Rhizophagus irregularis*. We performed an ultra-low input RNA-seq analysis, SMART-seq2, comparing five extraradical structures, germ tubes, runner hyphae, branched absorbing structures, immature spores, and mature spores. In addition, we reanalyzed the recently reported RNA-seq data comparing intraradical hyphae and arbuscules. Our analyses captured the distinct features of each structure and revealed the structure-specific expression patterns of genes related to absorption, metabolism, and transfer of nutrients. Of note, the transcriptional profiles suggest the distinct functions of branched absorbing structures in nutrient absorption. These findings provide a comprehensive dataset to advance our understanding of the transcriptional dynamics of fungal nutrition in this symbiotic system.

## Introduction

Arbuscular mycorrhizal fungi (AMF) establish symbiotic relationships with most terrestrial plants ^1^. The central function of these relationships is nutrient exchange, in which AMF supply plants with mineral nutrients and water, in exchange for carbon sources ^2^. Through this trade, AMF play important roles in terrestrial ecosystems ^3, 4^, and contribute to an increase in agricultural productivity ^5^. Therefore, it is important to understand the regulation of absorption, metabolism and transfer of nutrients associated with the symbiotic state of AMF.

Many studies to date have shown the regulation of absorption, metabolism and transfer of nutrients in AMF and the genes involved in these processes. Some of these studies investigated the differences between developmental stages, such as between germinating spores, intraradical mycelium (IRM), and extraradical mycelium (ERM). However, the differences between the structures comprising each developmental stage are not well understood. In pre-symbiotic stages, AMF grow curved, thin hyphae called germ tubes (GT) from spores (Fig. 1a). When the GT attach to host roots, the AMF start to grow intraradical hyphae (IRH) and develop arbuscules (ARB). As the plant inorganic phosphate (Pi) transporters, ammonium transporters, and lipid metabolism enzymes required for arbuscular mycorrhizal (AM) symbiosis are specifically expressed in arbuscule-containing cells, it is thought that the transport of phosphorus, nitrogen and lipids occurs in arbuscules ^6–12^. The spatial expression patterns of the AMF genes required for nutrient transfer and metabolism, however, have not been well examined. Moreover, the site of saccharide transport is still under debate ^13, 14^. Although Zeng *et al.* (2018) recently reported the transcriptome comparing the IRH and the ARB, the expression patterns of genes putatively related to nutrient transfer and metabolism have not been analyzed in detail. On the other hand, AMF extend ERM in order to absorb nutrients from soil and produce spores. Straight thick hyphae called runner hyphae (RH) are spread out from the root (Fig. 1b, c), and subsequently, highly branched hyphae called branched absorbing structures (BAS) differentiate on the RH (Fig. 1b, c) ^15^. Daughter spores are formed on the BAS, and gradually mature (Fig. 1b, d) ^15^. The features of each extraradical structures, responsible for nutrient absorption and metabolism, are completely unknown. Previous studies speculated that the BAS have important roles for nutrient absorption because of their cytological similarity with ARB and because the pH of the media around the BAS forming spores decreases, presumably by the secretion of compounds for phosphate solubilization ^15, 16^. However, there is no direct evidence to show nutrient absorption by the BAS. Several studies have reported transcriptome analyses comparing the ERM with the IRM or germinating spores ^17–20^, but to our knowledge, there are no reports that comprehensively investigate the spatial gene expression pattern among extraradical structures.

**Fig. 1.**
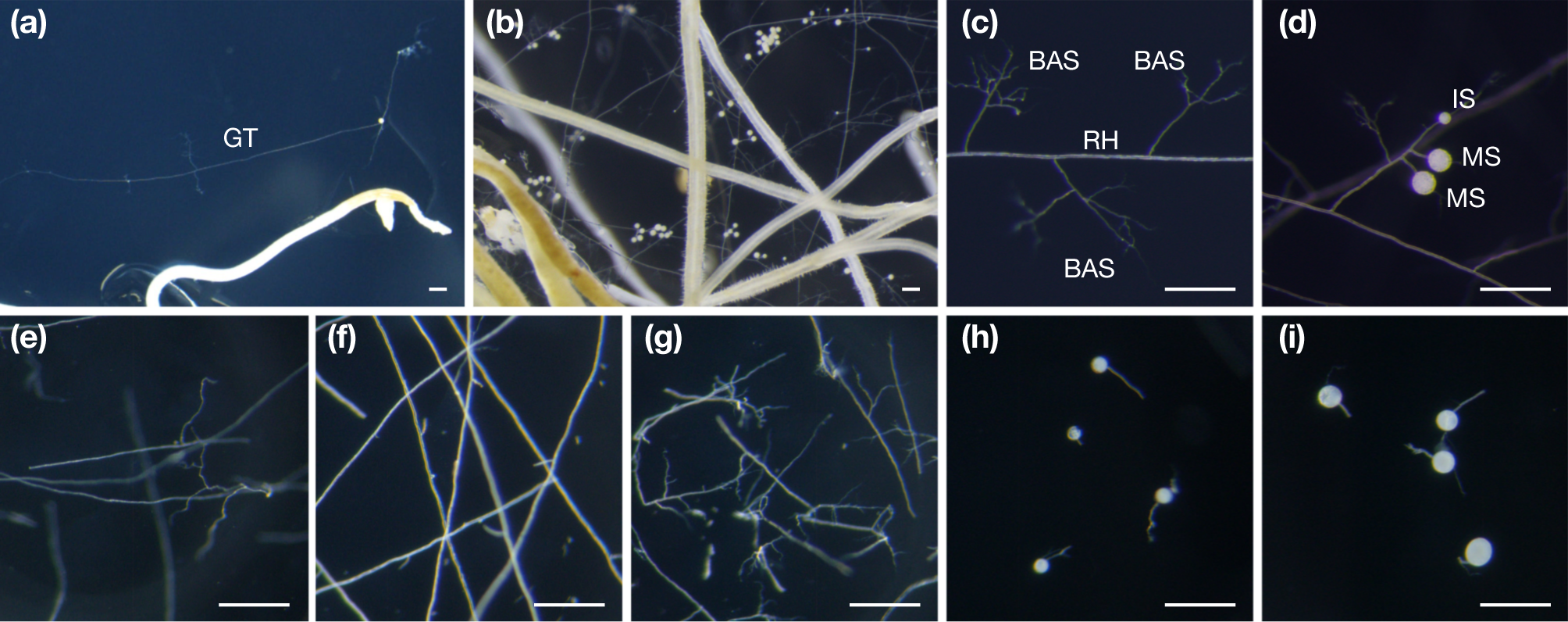
Extraradical structures. **(a)** 1-week-cultured *R. irregularis* in the pre-symbiotic stage. **(b–d)** 6-week-cultured *R. irregularis* in the symbiotic stage; global view **(b)**, runner hyphae and branched absorbing structures **(c)**, and immature and mature spores **(d)**. **(e–i)** Samples for RNA-seq analysis; germ tubes **(e)**, runner hyphae **(f)**, branched absorbing structures **(g)**, immature spores **(h)**, and mature spores **(i)**. Bar = 200 μm. GT, germ tubes; RH, runner hyphae; BAS, branched absorbing structures; IS, immature spores; MS, mature spores.

Here, we show the structure-specific transcriptome of the model AMF species, *Rhizophagus irregularis*. We individually collected five extraradical structures, GT, RH, BAS, immature spores (IS), and mature spores (MS), and performed RNA-seq analysis. To overcome the low volume of sample available, we applied the SMART-seq2 method, in which we were able to construct RNA-seq libraries from as little as 10 pg of RNA ^21, 22^. In addition, we reanalyzed the most recent RNA-seq data comparing ARB and IRH ^23^, focusing on the genes possibly involved in nutrient transfer to host plants. Our analyses captured the distinct features of each structure and revealed structure-specific expression patterns of genes related to absorption, metabolism, and transfer of nutrients. These results provide a basic dataset to understand nutrient exchange, which is the central function of AM symbiosis.

## Materials and Methods

### Biological materials and growth conditions

Hairy roots of carrot (*Daucus carota*) generated in Tsuzuki *et al.* (2016) were grown on M medium (731 mg l^−1^ MgSO_4_·7H_2_O, 80 mg l^−1^ KNO_3_, 65 mg l^−1^ KCl, 4.8 mg l^−1^ KH_2_PO_4_, 288 mg l^−1^ Ca(NO_3_)_2_·4H_2_O, 10 g l^−1^ Sucrose, 8 mg l^−1^ NeFeEDTA, 0.75 mg l^−1^ KI, 6 mg l^−1^ MnCl_2_·4H_2_O, 2.65 mg l^−1^ ZnSO_4_·7H_2_O, 1.5 mg l^−1^ H_3_BO_3_, 0.13 mg l^−1^ CuSO_4_·5H_2_O, 0.0024 mg l^−1^ Na_2_MoO_4_·2H_2_O, 3 mg l^−1^ glycine, 0.1 mg l^−1^ thiamine hydrochloride, 0.1 mg l^−1^ pyridoxine hydrochloride, 0.5 mg l^−1^ nicotinic acid, 50 mg l^−1^ myo inositol, 4 g l^−1^ gelrite) ^24^ at 28°C in the dark. Monoxenic cultures of the hairy root and *Rhizophagus irregularis* DAOM 197198 (Premier Tech, Rivière-du-Loup, Canada) were performed as described in Tsuzuki *et al.* (2016).

### Sample preparation and RNA extraction

Extraradical structures were dissected by surgical knife under stereoscopic microscope. To avoid wounding stress responses, dissected structures were frozen within *ca.* 1 min in liquid nitrogen. After smashed in RNA extraction buffer, enough amount of structures for a replicate were gathered into a tube. GT were sampled from 7 days after inoculation (dai) monoxenic culture plates. The hyphae which did not attach to the hairy root from 10–30 germinating spores were collected for a replicate. We collected 20–50 BAS which do not contain spores, 5–10 RH (ten BAS were dissected from each RH), five IS, and five MS for a replicate from 42 dai plates. As RH and BAS were dissected at the base part of BAS, BAS samples did not include RH, while RH include small amount of BAS. Spores smaller than 100 μM in diameter were defined as immature spores, while the larger ones were defined as mature spores. RNA was extracted using NucleoSpin RNA XS (Macherey-Nagel, Düren, Germany). As the amount of RNA extracted from spores were often too much to amplify as for the same cycles as hyphal samples, 2–100% of extracted RNA was used for later steps. The amount of samples used to synthesize cDNA were summarized in Table S1.

### Library synthesis and sequencing

RNA was reverse-transcribed and amplified using the SMART-Seq v4 Ultra Low Input RNA Kit for Sequencing (Clontech, Mountain View, CA, USA) ^21, 22^. The quality and the quantity of cDNA was checked using the Agilent 2100 Bioanalyzer with the High Sensitivity DNA kit (Agilent Technologies, Santa Clara, CA, USA). cDNA was sheared using the COVARIS S2 system (Covaris Inc., Woburn, MA, USA) and converted to RNA-seq libraries using the Low input library prep kit (Clontech, Mountain View, CA, USA). Each library was diluted to 2 nM. Four replicates were prepared for each structure. The amount of cDNA and libraries were summarized in Table S1. 102 bp single-end reads were obtained using the HiSeq 1500 (Illumina, San Diego, CA, USA). The RNA-seq data comparing the IRH and ARB were obtained from BioProject PRJNA389248.

### Differentially expressed gene (DEG) analysis

These reads were trimmed using Trimmomatic (v0.33) ^25^. Trimmed reads were mapped to the latest genome data ^26^ by using tophat2 (v2.1.0) ^27^. The read counts were calculated by HT-seq (v0.6.1) ^28^. Information about raw reads, trimming, and mapping were summarized in Table S2. DEGs (|log_2_ FC| > 1, FDR < 0.05) were detected using EdgeR-robust (v3.18.1) ^29, 30^. To compare the expression profiles among structures, count per million (CPM) of each gene was normalized by trimmed-mean-of-M-values (TMM) using edgeR ^29^, and subsequently scaled using the scale function in R (v3.4.1). PCA was performed using the prcomp function in R, using all expressed genes. Non-expressed genes were eliminated from this analysis. For PCA comparing the DEGs, the expression levels of DEGs in each structure calculated by EdgeR-robust were scaled using the scale function in R and PCA was performed using the prcomp function in R. K-means classification of DEGs was performed using the kmeans function in R. Gene ontology was annotated by Blast2GO (v4.1) ^31^. Enrichment analyses were performed by goseq ^32^.

## Results and discussion

### Structure-specific transcriptome analysis of *R. irregularis* using the low input RNA-seq method

To investigate the physiological features of extraradical structures of AMF, we performed transcriptome analysis in *R. irregularis* DAOM197198, comparing five extraradical structures, GT, RH, BAS, IS, and MS (Fig. 1e–i). Although RH, IS, and MS samples contain small amount of BAS (Fig. 1f,h,i), the influence of the contamination seems negligible, because the amount of RNA extracted from a BAS was much fewer than that from other structures (Table S1). As it was difficult to collect sufficient amounts of samples for the regular RNA-seq method, we constructed the RNA-seq libraries using the SMART-seq2 method, a low input library preparation method ^21, 22^. The libraries were sequenced using the Illumina HiSeq1500 and the reads obtained were analyzed based on the latest genome data ^26^ (Table S2).

We first validated the sampling and sequencing method by using principal component analysis (PCA) of gene expression profiles among structures and replicates (Fig. 2a). PC1 clearly discriminated between hyphae and spores, while PC2 captured the maturation of spores. Although hyphae samples were not clearly separated in the PC1-PC2 plane, they were separated in the PC3-PC4 plane. These results indicated that the low input RNA-seq method successfully detected the differences in gene expression patterns among the extraradical structures.

**Fig. 2.**
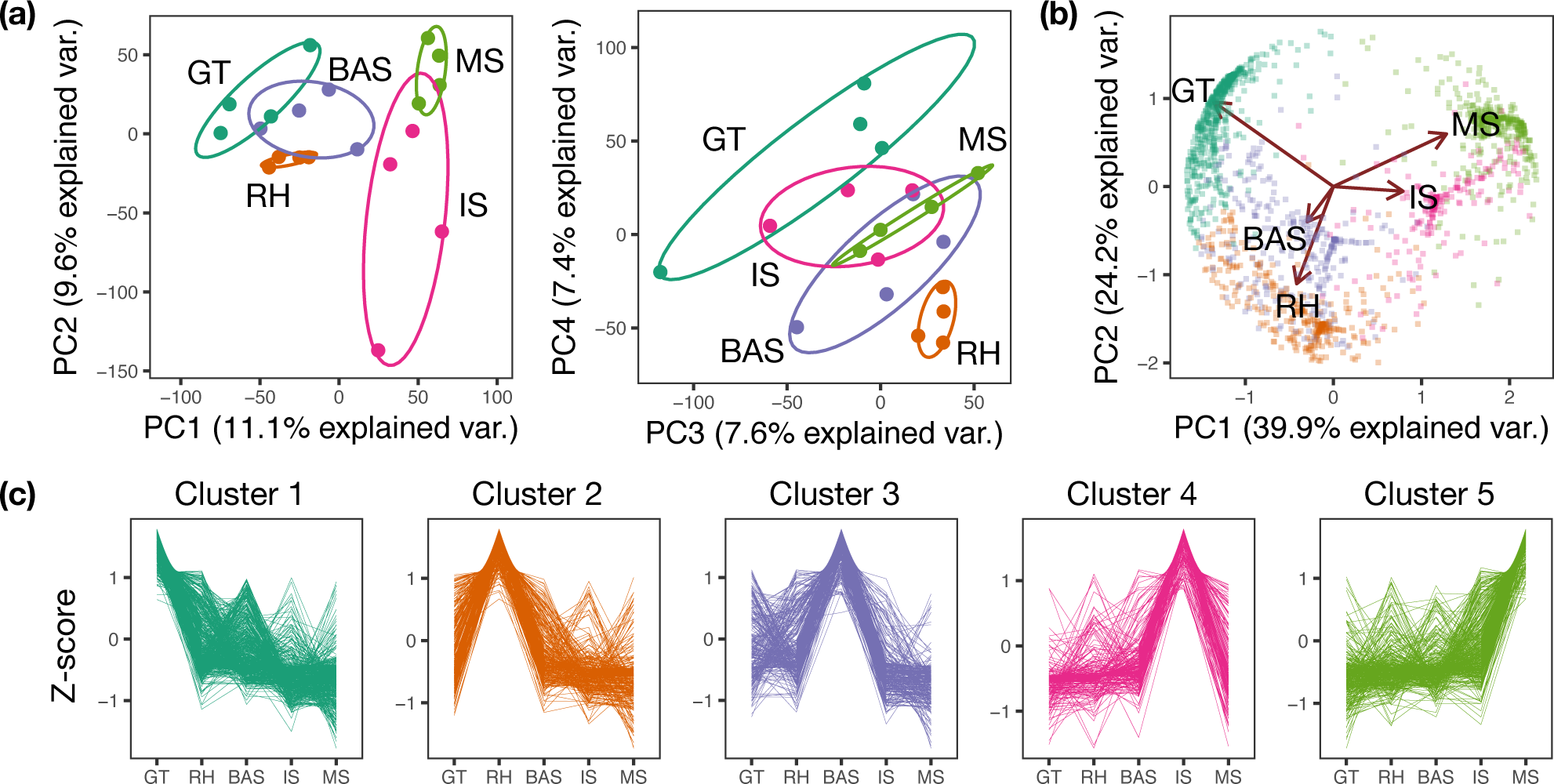
Detection of structure-specific gene expression. **(a)** Principal component analysis (PCA) of gene expression profiles between structures and replicates. Ellipses represent a 68 % confidence region for each structure. **(b)** PCA analysis of 1,977 differentially expressed genes (DEGs). DEGs were classified into five clusters using the K-means method. **(c)** The scaled expression levels of genes in each cluster. Genes in the same clusters are labelled in the same color. GT, germ tubes; RH, runner hyphae; BAS, branched absorbing structures; IS, immature spores; MS, mature spores.

Among the 19,918 genes expressed in all the samples, 1,977 genes were differentially expressed among the extraradical structures (Table S3). To visualize the expression pattern of the differentially expressed genes (DEGs), we again performed PCA. The axes of the five structures indicated that PC1 demonstrates whether the DEGs were expressed in hyphae or spores and that PC2 demonstrates whether they were expressed in pre-symbiotic hyphae, GT, or symbiotic extraradical hyphae, RH and BAS (Fig. 2b). This result seems reasonable considering the functional similarity of the structures.

DEGs were densely mapped near the axis of each structure, showing that many DEGs were highly expressed in one structure only (Fig. 2b). Therefore, we classified the DEGs into five clusters using the K-means method. Clusters 1, 2, 3, 4 and 5 were represented by genes highly expressed in the GT, RH, BAS, IS, or MS, respectively (Fig. 2b, c). To investigate the function of genes in each cluster, we performed gene ontology (GO) enrichment analysis using the GO term Biological Process (Table S4). In cluster 3, the GO term ‘transmembrane transport’ (GO:0055085) was enriched. Although called ‘branching absorbing structures’, there is no direct evidence to show nutrient absorption by the BAS ^15, 16^. Our results, however, strongly suggest the involvement of the BAS in nutrient absorption. The enrichment of many GO terms involved in DNA replication and nuclear cell division in cluster 4 appears to reflect the burst in mitosis that occurs during spore maturation ^33^. Thus, the results of the GO enrichment analysis are consistent with previous observations, suggesting that our RNA-seq analysis captured the characteristic transcriptional profiles of each structure.

### Survey of genes related to absorption, metabolism and transfer of nutrients

To date, many studies have identified the genes related to absorption, metabolism, and transfer of nutrients in *R. irregularis*. These genes, however, have been annotated based on past transcriptome or genome data and are sometimes named differently in different reports (e.g., *GiPT* in Maldonado-Mendoza et al., 2001 and *RiPT1* in Walder et al., 2016). Therefore, we re-annotated these genes based on our current genome data ^26^, and re-named them uniformly, using the reported gene names (Table S5). In addition, we searched unidentified genes, putatively related to absorption, metabolism, and transfer of nutrients, based on orthology with the genes of *Saccharomyces cerevisiae* or *Aspergillus nidurans*, as detected by the OrthoFinder software ^26, 34^ (Table S5). Next, we examined the expression patterns of these genes, as discussed below.

### Absorption and metabolism of mineral nutrients

Phosphorus and nitrogen are important nutrients absorbed by AMF from soil and then transferred to host plants ^35, 36^. AMF absorb Pi from soil using two types of Pi importer, the H^+^ Pi symporter and –Pi symporter. AMF also utilize organic phosphate by secreting acid phosphatase. Most of the Pi absorbed by the ERM is transformed into polyphosphate, presumably in the tonoplast by the vacuolar transporter chaperone (VTC) complex, and then transported to the IRM through vacuoles ^41–44^. Polyphosphate is probably degraded to monophosphate by polyphosphatase and subsequently released from AMF ^45^. Although the mechanism by which AMF release Pi is largely unknown, orthologs of Suppressor of Yeast Gpa1 (SYG1), which share domains with plant and animal Pi exporters, are strong candidates for the Pi exporter ^46, 47^. In the case of nitrogen, AMF absorb NO_3_^−^ and NH ^+^, and transform them into arginine in the ERM. Arginine is transported into the IRM through vacuoles and degraded into NH_4_^+^ before being transferred to host plants ^36, 48–50^. Many genes which encode transporters and enzymes involved in these processes have been identified ^49, 51–53^.

Two of three genes encoding H^+^–Pi co-transporters, *PT1* (*GiPT* in Maldonado-Mendoza et al., 2001 and *RiPT1* in Walder et al., 2016) and *PT2*, were differentially expressed among the extraradical structures, while *PT3* and two genes encoding Na^+^–Pi co-transporters, *PT5* and *PT6*, were not differentially expressed and showed relatively low expression levels (Fig. 3a, Table S5). *PT1* was the most highly expressed gene among the Pi transporters and showed the highest expression level in the BAS (Fig. 3a). Expression of *ACP1*, a gene encoding extracellular acid phosphatase, was increased with the formation of immature spores from BAS, and reached the maximum in MS (Fig. 3b). This correlates with previous observations that the pH of the medium around the BAS-containing spore has decreased ^16^. These results suggest the importance of the BAS for Pi absorption. *VTC1*, which is thought to be involved in Pi polymerization, was highly expressed in the RH, while *PPN3*, a gene encoding endopolyphosphatase, was highly expressed in the BAS, suggesting that Pi metabolism is specifically regulated in the RH and BAS (Fig. 3c,d). The expression levels of *SYG1* orthologs were low in the extraradical structures (Table S5).

**Fig. 3.**
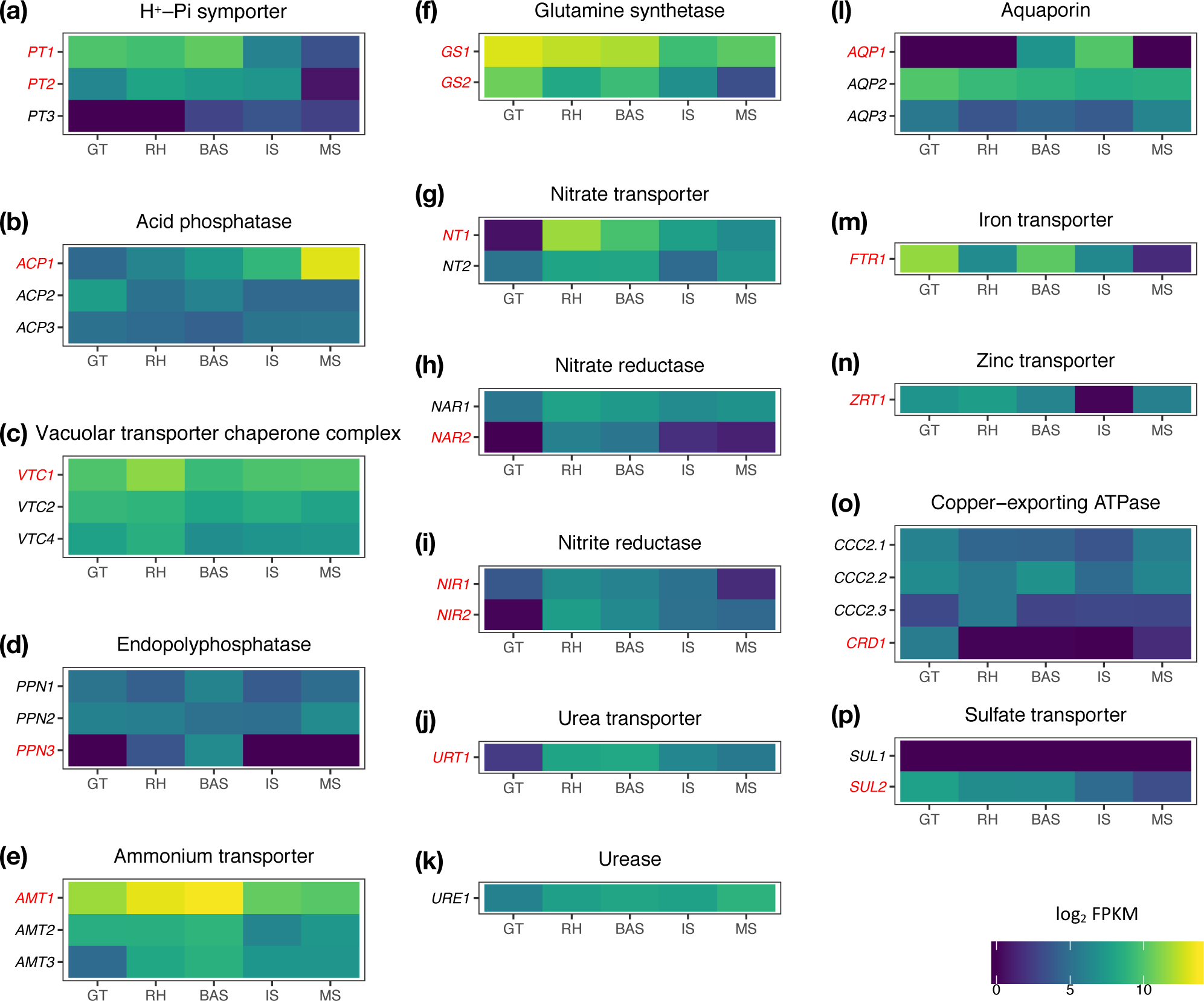
Expression patterns of genes related to mineral nutrient absorption and metabolism. Log_2_ fragments per kilobase of exon per million reads mapped (FPKM) in each structure of the genes encoding H^+^ Pi symporter **(a)**, acid phosphatase **(b)**, vacuolar transporter chaperone complex **(c)**, endophosphatase **(d)**, ammonium transporter **(e)**, glutamine synthase **(f)**, nitrate transporter **(g)**, nitrate reductase **(h)**, nitrite reductase **(i)**, urea transporter **(j)**, urease **(k)**, aquaporin **(l)**, iron transporter **(m)**, zinc transporter **(n)**, copper-exporting ATPase **(o)**, and sulfate transporter **(p)**. The expression of genes in red letters was statistically significantly different among the structures (FDR < 0.05). GT, germ tubes; RH, runner hyphae; BAS, branched absorbing structures; IS, immature spores; MS, mature spores.

The expression of *AMT1* was dominant in the extraradical structures among ammonium transporters (Fig. 3e). *AMT1* was expressed at relatively high levels in all structures examined, especially in the BAS (Fig. 3e). The expression of genes encoding glutamine synthase, the first step in ammonium assimilation, was highest in the GT (Fig. 3f). Considering the expression pattern of these genes, AMF may use ammonium as a nitrogen source in all life stages. Conversely, genes encoding nitrate transporter, nitrate reductase, and nitrite reductase were highly expressed in the RH while strongly suppressed in the GT (Fig. 3g–i), indicating that the RH has a central role in nitrate metabolism and that *R. irregularis* cannot utilize nitrate in pre-symbiotic stages. As the reduction of nitrate to ammonium requires high energy ^54^, it seems reasonable that the AMF assimilates nitrate in the symbiotic stages only, where they can obtain sufficient carbon sources from host plants. The expression patterns of the genes encoding urea transporter and urease were similar to those of nitrate-related genes (Fig. 3j,k). Some arginine metabolism genes were differentially expressed among the extraradical structures, but their expression did not follow a particular pattern (Table S5).

One of three genes encoding aquaporin, *AQP1*, was expressed specifically in the IS (Fig. 3l). As AQY1, a yeast spore-specific aquaporin, may function in spore maturation ^55^, *AQP1* might also be involved in this process. Genes encoding iron transporter, zinc transporter, copper-exporting ATPase, and sulfate transporter were also differentially expressed among the extraradical structures (Fig. 3m–p).

### Carbon absorption and metabolism

AMF strongly depend on plants as their source of carbon ^2, 56^, although they have the ability to absorb acetate and monosaccharide on the ERM and germinating spores ^13, 56, 57^. AMF acquire monosaccharides by using monosaccharide transporters (MSTs) ^13, 58, 59^, and catabolize them via the glycolytic pathway and TCA cycle ^60, 61^. On the other hand, they synthesize trehalose and glycogen to maintain monosaccharide concentrations inside the cells and to transport carbon to the ERM ^56, 62, 63^. In addition, recent studies have revealed direct lipid transfer from plants to AMF ^6, 7, 64^. AMF can elongate and desaturate fatty acids but not synthesize them ^65^, because they lack cytosolic fatty acid biosynthesis genes ^26, 66–69^, indicating that all fatty acids in AMF are plant-derived. The molecular form of transferred lipids is still unknown, but the predicted catalytic properties of the plant enzymes required just before lipid transfer imply that *sn-*2-palmitoylglycerol is a strong candidate ^6–8, 64, 70, 71^. AMF produce ATP and synthesize saccharides from fatty acids via the β-oxidation pathway, the glyoxylate cycle, and the gluconeogenesis pathway ^57, 58, 67, 72^.

Consistent with the previous observation that AMF can absorb acetate and monosaccharide from the ERM ^13, 57^, the expression of acetate transporters, carboxylate transporters and monosaccharide transporters was clearly detected in the extraradical structures (Fig. 4a–c). As our previous phylogenetic analysis classified *RiMST6* ^59^ not into the orthogroup of monosaccharide transporters (OG0000019) but into that of carboxylate transporters (OG0001862) ^26^, we re-named it *JEN2* (Fig. 4a,b and Table S5). MST2 is a monosaccharide transporter which is expressed in the IRM and required for mycorrhiza formation involving arbuscule development ^13^. While expression of *MST2* in the ERM was not detected previously ^13^, a relatively low but clear level of expression was observed in the BAS (Fig. 4a). In the extraradical structures, *MST4* and *MST5* showed relatively high expression levels among *MST*s (Fig. 4a), suggesting the contribution of these genes to monosaccharide absorption in ERM. *ACT2* and *ACT3*, genes encoding acetate transporter, were expressed in a broad range of the extraradical structures (Fig. 4b). On the other hand, the expression of *JEN3* and *JEN4*, genes encoding carboxylate transporters, were significantly highly expressed in BAS (Fig. 4c) suggesting the importance of BAS for carboxylate absorption.

**Fig. 4.**
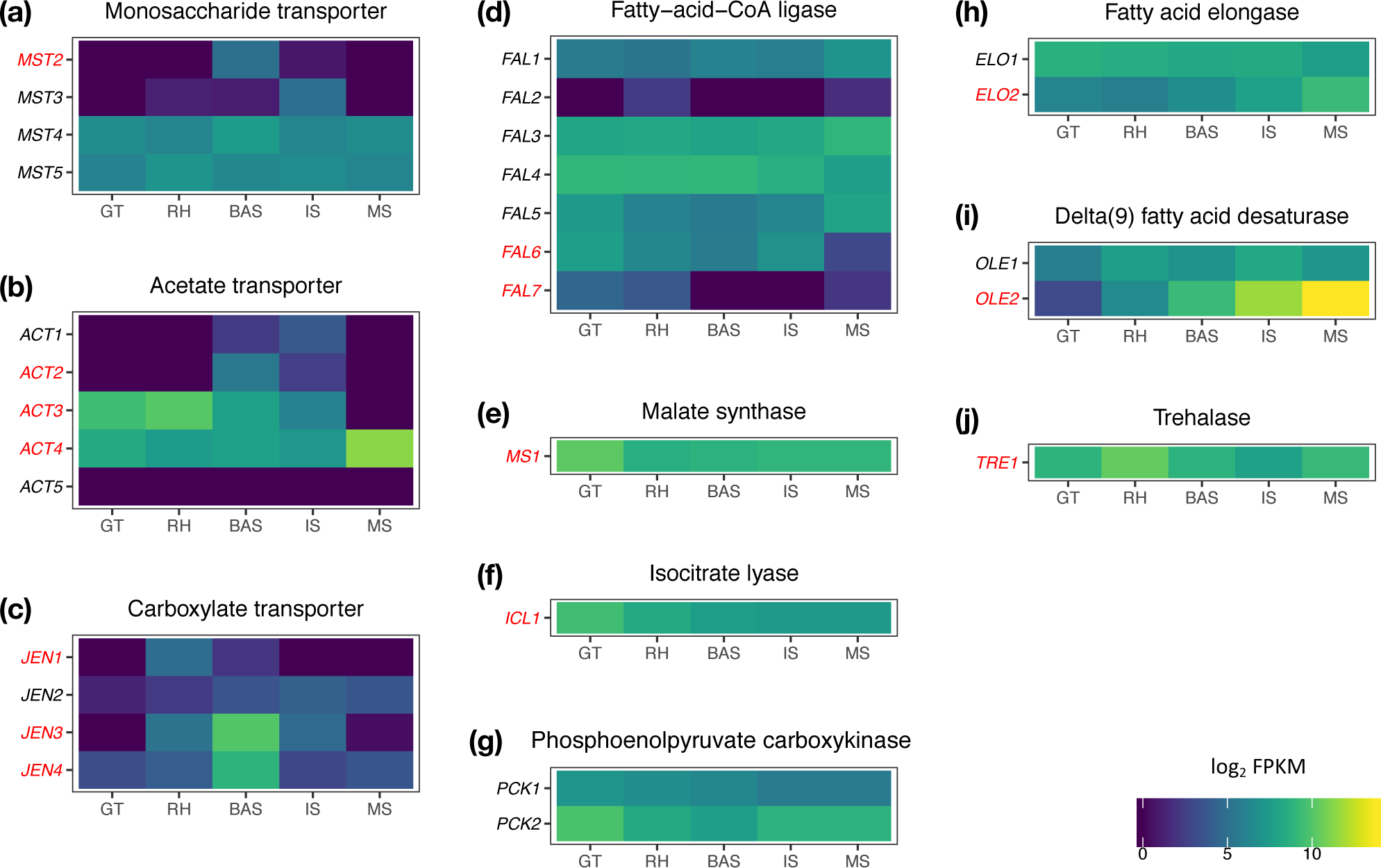
Expression patterns of genes related to carbon absorption and metabolism. Log_2_ fragments per kilobase of exon per million reads mapped (FPKM) in each structure of the genes encoding monosaccharide transporter **(a)**, acetate transporter **(b)**, carboxylate transporter **(c)**, Fatty-acid-CoA ligase **(d)**, malate synthase **(e)**, isocitrate lyase **(f)**, phosphoenolpyruvate carboxykinase **(g)**, fatty acid elongase **(h)**, delta 9-fatty acid desaturase **(i)**, and trehalase **(j).** The expression of genes in red letters was statistically significantly different among the structures (FDR < 0.05). GT, germ tubes; RH, runner hyphae; BAS, branched absorbing structures; IS, immature spores; MS, mature spores.

Two of seven genes encoding fatty-acid-CoA ligase, *FAL6* and *FAL7*, which functions in the β-oxidation pathway, were differentially expressed with statistical significance, and showed the highest expression levels in the GT (Fig. 4d). In addition, both of the genes encoding glyoxylate cycle-specific enzymes, isocitrate lyase and malate synthase, were significantly highly expressed in the GT (Fig. 4e,f). Genes encoding phosphoenolpyruvate carboxykinase, an enzyme functioning in gluconeogenesis, were also highly expressed in the GT, although the differences were not statistically significant (Fig. 4g). These expression patterns are consistent with reports that most of the carbon storage in the spores of AMF is in the form of lipids and fatty acids ^73, 74^. Conversely, two genes encoding enzymes for the modification of fatty acids, fatty acid elongase and delta 9-fatty acid desaturase, were highly expressed in the MS (Fig. 4h,i). These genes might have an important contribution to fatty acid storage in spores. On the other hand, the gene encoding trehalase was highly expressed in the RH (Fig. 4j), presumably reflecting the abundant saccharides available from host plants in the symbiotic stages.

### Nutrient transfer and metabolism in the intraradical phase

Although Zeng et al. (2018) provided high quality RNA-seq data comparing the IRH and ARB, genes related to nutrient transfer and metabolism have not been analyzed in detail. To investigate the function of the IRH and ARB in nutrient exchange between AMF and host plants, we reanalyzed the RNA-seq data. PCA between samples clearly discriminated between the IRH and ARB samples (Fig. 5a).

**Fig. 5.**
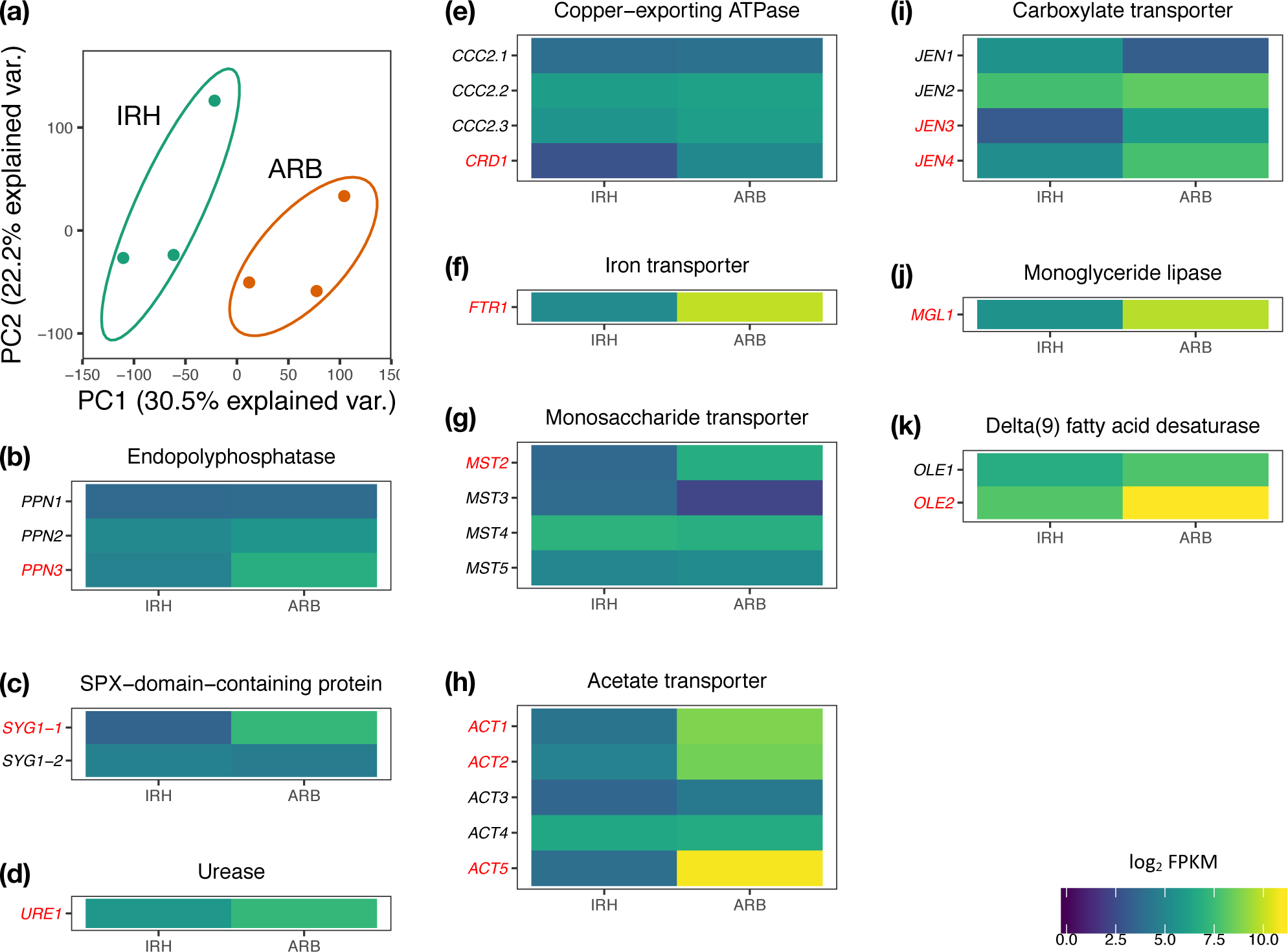
Differentially expressed genes between intraradical hyphae and arbuscules. **(a)** Principal component analysis (PCA) of gene expression profiles of intraradical hyphae and arbuscule samples. IRH, intraradical hyphae; ARB, arbuscule. **(b–k)** Log_2_ fragments per kilobase of exon per million reads mapped (FPKM) in intraradical hyphae or arbuscules of the genes encoding endophosphatase **(b)**, SPX-domain-containing protein **(c)**, urease **(d)**, copper-exporting ATPase **(e)**, iron transporter **(f)**, acetate transporter **(g)**, monosaccharide transporter **(h)**, carboxylate transporter **(i)**, Monoglyceride lipase **(j)**, and delta 9-fatty acid desaturase **(k)**. The expression of genes in red letters was statistically significantly different between the IRH and ARB (FDR < 0.05).

We then analyzed DEGs between the IRH and ARB. Among the phosphate-related genes examined (Table S5), *PPN3*, a gene encoding endophosphatase, and *SYG1-1*, a gene encoding the homolog of a Pi exporter, were mainly expressed in the ARB compared to the IRH (Fig. 5 b,c). Although the functions of these genes have not been investigated, it is possible that they are involved in Pi release from AMF. In concordance with the report that arginine is degraded into ammonium before nitrogen is transferred to host plants ^50^, urease, which catalyzes the conversion of urea to ammonium in the final step of arginine degradation, was highly expressed in the ARB (Fig. 5d). *CRD1*, a gene encoding copper-exporting ATPase, and *FTR1*, a gene encoding an iron transporter, were also highly expressed in the ARB (Fig. 5e,f), indicating the function of ARB for copper and iron exchange between plants and AMF. Although *MST2* was expressed in both the IRH and ARB, its expression level was higher in the ARB (Fig. 5g), indicating that the ARB has a significant role in saccharide absorption. While *ACT1*, *ACT2*, and *ACT3* showed lower expression levels in extraradical structures among genes encoding acetate transporter (Fig. 4b), their expression levels were significantly high in ARB (Fig. 5h). On the other hand, two carboxylate transporters, *JEN3* and *JEN4*, which were significantly expressed in BAS (Fig. 4c), also showed higher expression levels in ARB compared to IRH with statistical significance (Fig. 5i). The expression of some genes encoding acetate transporters and carboxylate transporters was also induced in the ARB (Fig. 5h,i). Two lipid metabolism-related genes *MGL1* and *OLE2*, which encode monoacylglyceride lipase and delta 9-fatty acid desaturase, respectively, were highly expressed in the ARB (Fig. 5j,k). This result led us to hypothesize that *sn-*2-palmitoylglycerol is hydrolyzed into palmitoic acid and glycerol by MGL1, and palmitoic acid is desaturated by OLE2. Thus, analysis of the expression patterns of putative nutrient transfer genes has provided insight into the functions of the ARB for nutrient exchange between AMF and host plants.

## Concluding remarks

In this study, we performed transcriptome analysis comparing extraradical structures of the AMF *R. irregularis* to investigate the functions of these structures in nutrient absorption and metabolism. Our analysis generated structure-specific transcriptional profiles and detected DEGs among the structures. Further analysis of genes related to nutrient absorption and metabolism, and reannotation using the latest genome data, revealed the characteristic functions of each structure; for example, fatty acid consumption in the GT (Fig. 4d–g), nitrate absorption and reduction in the RH (Fig. 3f–i), Pi, ammonium, and carboxylate absorption in the BAS (Fig. 3a,e; Fig. 4c), and fatty acid modification in the MS (Fig. 4h,i). In addition, we reanalyzed the RNA-seq data comparing the IRH and ARB ^23^ and detected the expression of putative nutrient transfer genes in the ARB (Fig. 5). Although many studies have revealed the regulation of absorption, metabolism, and transfer of nutrients by AMF, the spatial regulations among the structures have remained largely unknown. Our data, therefore, provides comprehensive information for understanding the dynamics of nutrient regulation in this symbiotic system.

Of note, our results suggest distinctive functions between the RH and BAS for nutrient absorption and metabolism. Although they have morphological differences, it was not clear previously as to whether the RH and BAS have different transcriptional profiles, because they are not separated by either a cell membrane nor a cell wall and share a common cytoplasm ^15^. This question had not been approached previously, probably because of technical difficulties. Our RNA-seq analysis using the SMART-seq2 method clearly illustrated the difference in the transcriptome between the two structures. Although many genes related to nutrient absorption and metabolism were expressed in both structure, they showed distinctive expression patterns, indicating the importance of the RH for the absorption and reduction of nitrate (Fig. 3g–i) and that of the BAS for the absorption of Pi, ammonium, and carboxylate (Fig. 3a,e; Fig. 4c). These results provide a new perspective for investigating nutrient absorption and metabolism in AMF.

Environmental response of genes related to nutrients is an important aspect to understand the nutrition of the symbiotic system. It is plausible that the expression patterns of genes related to nutrients show different expression patterns from our results in response to nutrient conditions or other environmental factors in nature. As we collected the extraradical structure samples in an *in vitro* culture system, where the composition of nutrients and other conditions were controlled, our data would be references to examine the environmental responses.

To further understand the physiological functions of each structure, functional analysis of DEGs is required. Host-induced gene silencing (HIGS) and virus-induced gene silencing (VIGS), which are able to silence the genes of AMF in the IRM by using dsRNA produced by the host plant or a virus to infect the host plant, respectively, ^13, 18, 75^, will be useful tools for the analysis of the putative nutrient transfer genes expressed in the ARB. However, no method to manipulate gene expression in extraradical structures currently exists. Establishment of new technologies for transformation, gene silencing in extraradical structures and mutagenesis is therefore necessary.

In addition to absorption and metabolism of nutrients, extraradical structures have important functions such as infection in host plants, interaction with associated bacteria, and spore formation, the genetics of which have been poorly investigated. Our data provide fundamental information which should assist in these investigations.

## Availability

Raw sequence data has been deposited in the DDBJ Sequence Read Archive (DRA) under BioProject PRJDB6136.

## Acknowledgements

We thank Dr. Syusaku Tsuzuki, Dr. Naoya Takeda, Dr. Katsuharu Saito, and Prof. Tatsuhiro Ezawa for their critical suggestions and Ms. Yuko Ogawa, Ms. Sachiko Tanakta, Ms. Asami Tokairin and Ms. Yumi Yoshinori for their experimental support. We appreciate Dr. Katsushi Yamaguchi, Prof. Shuji Shigenobu, Functional Genomics Facility and Data Integration and Analysis Facility, NIBB Core Research Facilities for their technical support. This work was supported by ACCEL (JPMJAC1403) from the Japan Science and Technology Agency.

---

**Table S1.** Summary of samples used for RNA-seq library construction.

**Table S2.** Summary of RNA-seq reads and preprocessing.

**Table S3.** List of differentially expressed genes.

**Table S4.** Enriched GO terms (Biological Process) in differentially expressed genes.

**Table S5.** Genes related to nutrient absorption, metabolism, and transfer.

